# Elevated Thalamic Blood Flow in Self-Limited Epilepsy with Centrotemporal Spikes

**DOI:** 10.1101/2025.09.19.677437

**Authors:** Niki Iasinovschi, Elizabeth Tong, Lucia Bicknell, Fiona Mitchell Baumer

## Abstract

Children with self-limited epilepsy syndrome with centrotemporal spikes (SeLECTS) exhibit altered thalamocortical connectivity, but whether thalamic function itself is abnormal remains unclear. We investigated whether thalamic blood flow, a marker of metabolism, differs between children with SeLECTS and controls, and examined the effects of spike distribution and antiseizure medications (ASMs) on thalamic perfusion. In this retrospective cohort study, we identified consecutive children with SeLECTS who underwent magnetic resonance imaging (MRI) for epilepsy evaluation (n = 44) and age- and sex-matched children who underwent MRI for non-epilepsy indications (n = 35). We quantified thalamic blood flow via manual segmentation of cerebral blood flow (CBF) sequences obtained from arterial spin labeling MRI. Clinical variables including sedation use during MRI, daily ASM use, and spike distribution (unilateral or bilateral) were extracted from medical records. Children with SeLECTS demonstrated elevated thalamic blood flow compared to controls, with the most pronounced differences in specific subgroups. Children with unilateral spikes showed the highest CBF, particularly in the thalamus contralateral to spike activity. ASM use significantly modulated thalamic blood flow: children taking oxcarbazepine showed the highest CBF, while those on levetiracetam had CBF similar to controls. Unmedicated children showed intermediate elevations. These findings demonstrate that elevated thalamic blood flow may be intrinsic to SeLECTS pathophysiology, with different ASMs producing distinct neurobiological effects. The differential medication effects may relate to their clinical efficacy and provide neurobiological rationale for treatment selection in this common childhood epilepsy syndrome.

## 1. Introduction

Self-limited epilepsy with centrotemporal spikes (SeLECTS) is the most common focal epilepsy of childhood characterized by the emergence of epileptiform activity and seizures during sleep. Two-thirds of children with SeLECTS have predominantly unilateral spikes from one motor cortex, whereas one-third have bilateral spikes (Wirrell, 1998). Though SeLECTS comprises about 15% of diagnosed pediatric epilepsy cases, the etiology of the disorder is unknown (Zucconi, 2011). Prior work has focused on investigating the anatomy (Garcia-Ramos et al., 2015) and connectivity (Besseling et al., 2013) of the sensorimotor cortices, with recent work showing alterations in both structural (Thorn et al., 2020) and functional (Kwon et al., 2022) connectivity between the thalamus and the sensorimotor cortices. The thalamus is critical for the generation of sleep architecture (Aime & Adamantidis, 2022). In SeLECTS, the frequency of spikes inversely correlates with the frequency of sleep spindles, suggesting that spikes co-opt thalamocortical sleep circuits (Kramer et al., 2021). These studies suggest that interactions between the thalamus and cortex are disrupted; however, investigation of intrinsic thalamic function has been limited.

An important measure of thalamic function is thalamic metabolism. Abnormal thalamic metabolism, as quantified by fluorodeoxyglucose positron emission tomography (FDG PET) imaging, is present in children with another, more severe epilepsy with sleep-potentiated spikes, developmental and epileptic encephalopathy with spike wave activation in sleep (D/EE-SWAS) (Agarwal et al., 2016). Due to radiation exposure, FDG PET is not justified for use in SeLECTS. Thalamic metabolism can be estimated without radiation using cerebral blood flow (CBF) images (Telischak et al., 2015; Venkat et al., 2016) obtained from arterial spin labeling (ASL) magnetic resonance imaging (MRI). CBF quantifies perfusion, capturing information about oxygen, glucose, and nutrient delivery (Chiacchiaretta et al., 2017). Blood flow increases to meet metabolic demands in activated regions of the brain, and thus elevated CBF suggests higher metabolic activity (Iadecola, 2017).

To examine whether thalamic blood flow is altered in SeLECTS, we conducted a retrospective cohort study comparing thalamic CBF obtained during clinical MRI in consecutively imaged children with SeLECTS to those of children who underwent MRI for non-epilepsy indications, excluding children with disorders known to alter CBF (e.g. migraine headaches). For children with SeLECTS, we reviewed electroencephalograms (EEGs) recorded near the time of MRI in order to classify spike activity as bilateral or unilateral, and if unilateral, to determine whether the left or right hemisphere was involved. Sixty percent of children with SeLECTS have unilateral spikes, originating in either the left or right hemisphere, while 40% have spikes emerging from both hemispheres, occurring either synchronously or asynchronously (Wirrell, 1998). We posited that thalamic blood flow increases as a result of spike-wave activity and thus would be higher in children with SeLECTS than controls and, in patients with unilateral spikes, higher in the hemisphere ipsilateral to spikes. We also examined the effects of different antiseizure medications (ASMs) on thalamic CBF, hypothesizing that sodium channel blockers would reduce thalamic CBF based on their known metabolic suppression effects in other epilepsy syndromes (Joo et al., 2006; Theodore et al., 1989), while levetiracetam might have different effects given its distinct mechanism of action and clinical efficacy profile in SeLECTS (Kanemura et al., 2018).

## 2. Methods

### 2.1 Sample Selection

The Stanford University Institutional Review Board approved this study. Using the Stanford Research Repository database, we queried the medical record using International Classification of Disease 10 (ICD-10) codes and keywords. We searched between 2010 and 2022 for children with a diagnosis of SeLECTS who underwent 3.0T brain MRI between the ages of 3 and 15 years. We manually reviewed records to confirm children met diagnostic criteria for SeLECTS (Specchio et al., 2022), including focal motor seizures or tonic clonic seizures observed out of sleep and centrotemporal spikes on EEG. We confirmed that the MRI was read as clinically normal and contained T2-weighted, ASL, and CBF sequences.

Clinical controls were identified by searching for children with ICD-10 codes for headache without migraine, ocular abnormalities, syncope, cholesteatoma, vomiting, cutaneous facial abnormality, suspected aneurysm, delayed milestones, Li-Fraumeni syndrome, and Von Hippel-Lindau syndrome. Children were included only if the clinical 3.0 T MRI was read as normal and they had the requisite T2-weighted, ASL, and CBF sequences. We excluded children imaged for migraine diagnosis, as migraine is associated with increased CBF (Fu et al., 2022). Subjects with Li-Fraumeni syndrome or Von Hippel-Lindau syndrome undergo frequent surveillance MRIs (Clarke et al., 2022) for tumors; MRIs included were restricted to those obtained at least 2 years before a tumor diagnosis. If a control had multiple candidate MRIs, the MRI that maximized age-matching with the SeLECTS group was chosen.

Exclusion criteria for both groups included history of birth at less than 35 weeks gestational age, abnormal brain MRI, other epilepsy syndromes, neurosurgery, severe brain injury, intellectual disability, and severe medical conditions (e.g., autoimmune disease, congenital heart malformation) that would be expected to alter brain activity.

### 2.2 Clinical Information

The following information was gathered via chart review: age at MRI, age of first seizure, sex, clinical indication for MRI, and use of sedation (propofol, midazolam, fentanyl, lorazepam) during imaging. Increasing age, specifically in the age range of our study, has been associated with both increasing (Forkert et al., 2016) and decreasing thalamic CBF (Biagi et al., 2007; Chiron et al., 1992). The relationship between sex and thalamic CBF has not been well described, but cerebral metabolic rate, as measured by FDG PET, in the left thalamus has been found to be higher in females (Shin et al., 2021). The relationship between sedation and CBF in children is also unclear. Propofol, the most common agent used for MRI in children at our institution, reduces thalamic CBF in adults (Saxena et al., 2019). Children who receive propofol also have lower CBF than unsedated children, but they are also almost always younger too (Forkert et al., 2016). For SeLECTS, we recorded whether the child was on a daily antiseizure medication (ASM) at the time of MRI and how long they had been on ASM; ASM may indicate a more severe epilepsy course and the impact of ASMs on CBF is not yet defined. We classified children as taking or not-taking a daily ASM, and since different ASM have different effects on spikes (Kanemura et al., 2018; Vendrame et al., 2007), we additionally classified them by which specific medication they were prescribed.

#### 2.2.1 Spike Laterality

Since 60% of children with SeLECTS have predominantly unilateral spikes and 40% have bilateral spikes affecting both hemispheres (Wirrell, 1998), we assessed spike laterality in all children with SeLECTS. We reviewed the EEG report performed temporally closest to the MRI to determine if children had spikes reported in one hemisphere only (unilateral, noting left or right hemisphere) or in both hemispheres (bilateral). We confirmed reported findings via manual review of tracings by a board-certified pediatric epileptologist (FMB). We then classified all SeLECTS thalamic CBF measurements relative to spike laterality as per EEG review (Fig 3a). Children with bilateral spikes were categorized as having two “bilateral” thalamic CBF measurements. Children with unilateral spikes were classified as having one “unilateral spikes: ipsilateral” and one “unilateral spikes: contralateral” CBF measurement. Control children were categorized as having two “control” thalami.

### 2.3 Magnetic resonance imaging specifications

All subjects had undergone brain imaging on General Electric (GE) MR scanners at 3.0T GE with an 8-channel head coil. Pseudocontinuous ASL MR imaging was performed using a pseudocontinuous labeling duration of 1500 ms, post-labeling delay of 1500 ms. The T2-weighted MR imaging was performed with a 4632 ms repetition time, 10.5 ms echo time, flip angle 111 degrees, 24 cm field of view, and 3 excitations.

### 2.4 Thalamic CBF measurements

Since CBF maps lacked the anatomic detail to define a thalamic region of interest directly, we created a thalamic mask using the T2-weighted images which we co-registered to the CBF file. To do this, two independent reviewers (NI and LB) were trained by a board-certified pediatric neuroradiologist (ET) to identify thalamic borders. Each scan was segmented twice, once by each reviewer. The paintbrush tool in ITK-SNAP (Yushkevich et al., 2006) was used to define the left and right thalamus separately on each axial slice (Figure 1) of the T2-weighted sequence, creating a 3-dimensional mask that captured the whole thalamus. Coronal and sagittal views were checked to ensure the quality of the mask.

**Figure 1.**
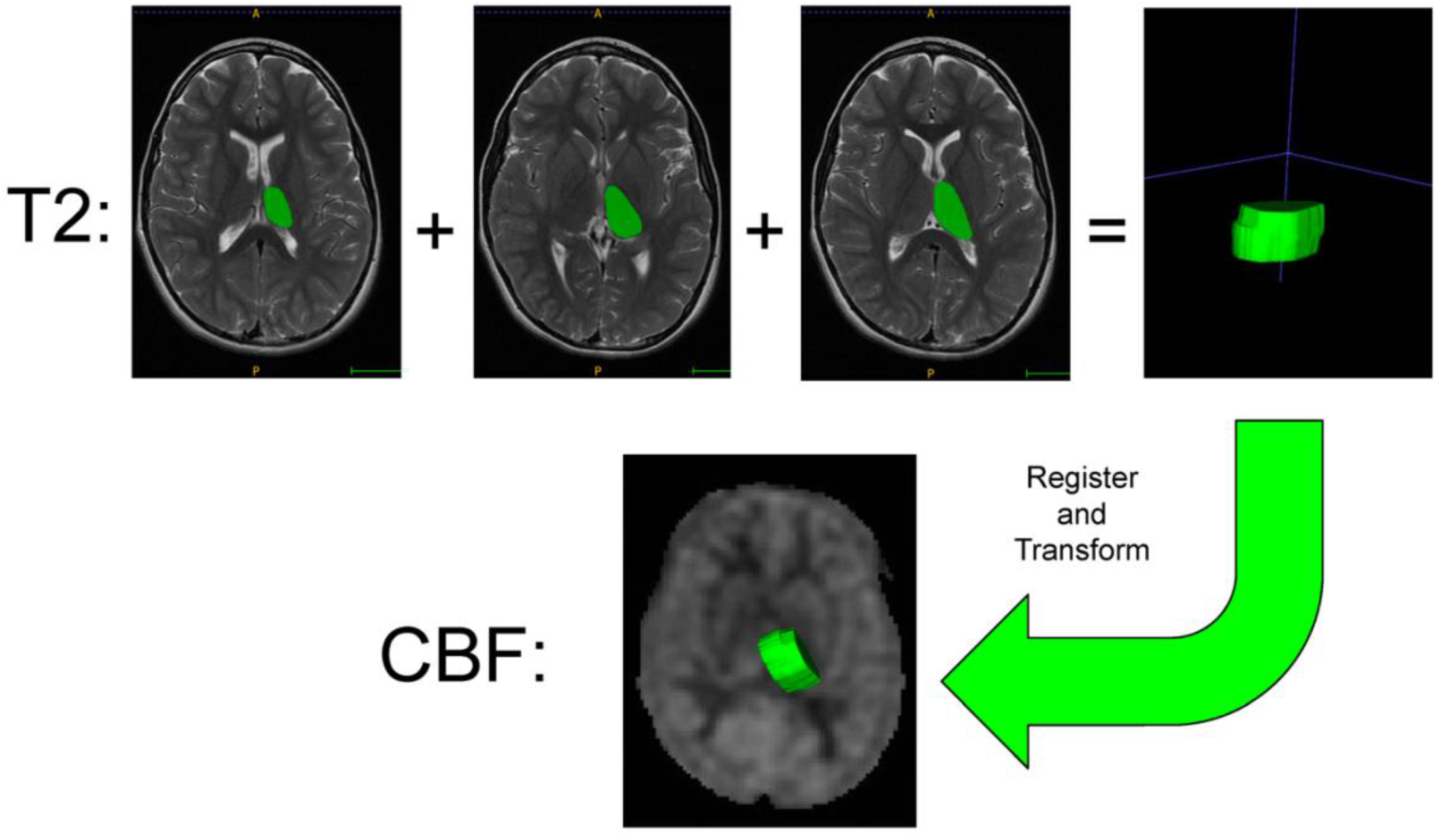
Segmentation of left thalamus on T2-weighted MRI and registration with CBF image. The left thalamus was segmented on the three consecutive axial slices of the T2-weighted MRI that displayed clear anatomical margins. In combination, the slices produced a mask of the thalamus. The CBF image was registered to the T2-weighted image, and the mask was used to obtain a CBF value for the left thalamus.

*Registration and CBF value extraction*. Thalamic masks were registered to the CBF map using a custom image-processing pipeline developed and implemented in MATLAB using the Insight Segmentation and Registration Toolkit. First, the CBF was registered to the corresponding T2-weighted sequence using rigid transformations. The affine transformation was determined using a linear interpolation and maximization of the mutual information metric. The calculated affine transformation was then used to initialize a b-spline transformation, for the fine nonlinear alignment of the thalamic masks to the ADC and CBF datasets. This b-spline transformation was optimized using linear interpolation and maximization of the mutual information metric. CBF values were extracted from the aligned thalamic regions. This process was completed for both raters’ masks, and the resulting values were averaged. Each subject had two CBF measurements: left thalamus and right thalamus.

All MRI sequences, T2-masks, and registration results were manually reviewed by a board-certified neuroradiologist (ET) to validate adequate quality for inclusion.

### 2.5 Statistical Analysis

Statistical analysis was performed using R Studio (R Core Team, 2023). Descriptive statistics for demographic variables of age and sex and medical variables of sedation, ASM, and spike laterality were calculated. Group differences in age, sex, and sedation use were analyzed using independent-sample t-tests and the Chi-square test for continuous and categorical values, as appropriate.

#### Group differences in thalamic CBF

We fit linear mixed models with thalamic CBF as the dependent variable and group (SeLECTS or control) as the independent variable of interest, with subject as a random intercept to account for repeated CBF measurements within each child (left and right thalamus) (West et al., 2014). We first assessed the group difference alone, then sequentially added potential confounders. We added age and sex to assess whether these variables attenuated the relationship between group and CBF, testing interaction terms to determine if group effects differed by age or sex. Finally, we included sedation status and a group-by-sedation interaction term to evaluate whether group differences were consistent across sedated and unsedated children, treating this as a sensitivity analysis to assess the robustness of our primary findings.

#### Spike Laterality

We first examined if there were any intrinsic differences in left versus right thalamic CBF by comparing these values in control participants using a paired t-test. To test if thalamic CBF was affected by location of spikes, we fit mixed models defining the group variable as one of four levels: Control; Bilateral spikes; Unilateral Spikes: Ipsilateral; Unilateral spikes: Contralateral), adjusting for age, sex, and sedation status.

#### Antiseizure Medications (ASMs)

To assess the impact of ASMs, we fit mixed models defining group as one of four levels (Control; SeLECTS not on ASM; SeLECTS on Oxcarbazepine; SeLECTS on Levetiracetam), again adjusting for age, sex, and sedation status.

Unstandardized effects (B) for the independent variables are reported with confidence intervals (CI) in our tables. From the models, we obtained estimated marginal means with standard error (SE) for CBF for each group and reported these in the text. We considered p<0.05 to be statistically significant.

## 3. Results

### 3.1 Subjects

We reviewed 311 consecutive charts of children identified by our search criteria as potentially having a SeLECTS diagnosis. Two hundred and twelve children were excluded because they did not meet diagnostic criteria for SeLECTS, and 19 were excluded due to a serious medical comorbidity. Of the 80 remaining children with SeLECTS, 44 had undergone an MRI containing a CBF sequence. We reviewed the charts of 795 potential controls to identify 35 controls who met the inclusion and exclusion criteria. Indications for control MRI scan were headache without migraine (n = 10), ocular abnormality (n = 7), cutaneous facial abnormality (n = 5), syncope (n = 3), Li-Fraumeni syndrome (n = 2), psychological issue (n = 2), Von Hippel-Lindau syndrome (n = 1), abnormal gait (n = 1), unknown growth disorder (n = 1), early puberty (n = 1), night sweats (n = 1), and sarcoma of foot with no metastasis (n = 1).

### 3.2 Demographic & Clinical Information (Table 1)

There were no significant group differences in age or in the relative proportion of males to females. Control children were more likely to have received sedation for the MRI. Of those receiving sedation, all received propofol except 3 children with SeLECTS (lorazepam n = 2, ketamine n = 1). More children with SeLECTS had unilateral spikes (12 left only, 12 right only) than bilateral spikes. Of the 28 children taking ASMs, 13 were on levetiracetam, 14 on oxcarbazepine, and one on lacosamide. The average duration of ASM use at time of MRI was 0.90 ± 1.6 years. SeLECTS MRIs occurred 0.97 ± 1.5 years after first seizure.

**Table 1.**
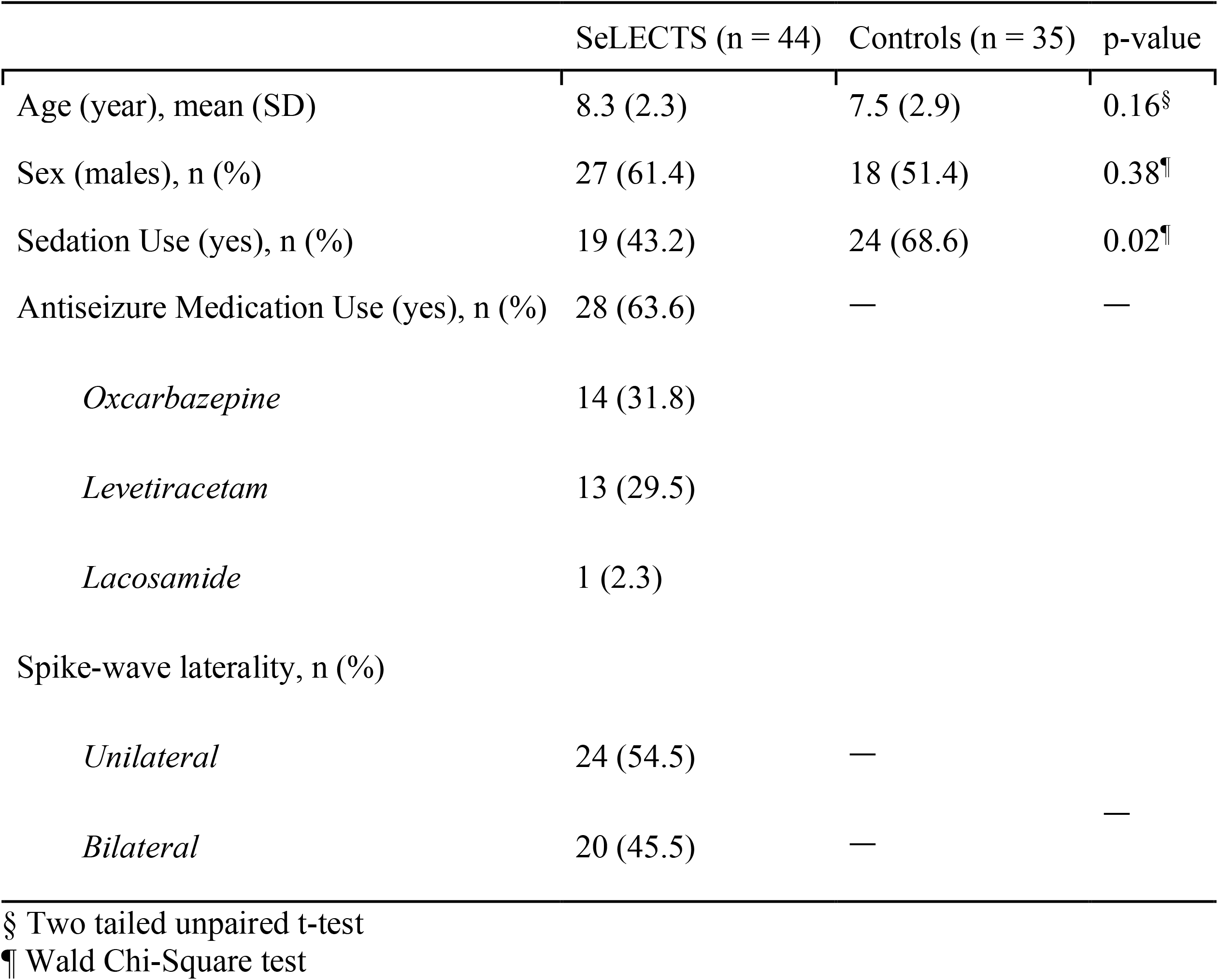
Demographic & Clinical Variables.

|  | SeLECTS (n = 44) | Controls (n = 35) | p-value |
| --- | --- | --- | --- |
| Age (year), mean (SD) | 8.3 (2.3) | 7.5 (2.9) | 0.16<sup>§</sup> |
| Sex (males), n (%) | 27 (61.4) | 18 (51.4) | 0.38<sup>¶</sup> |
| Sedation Use (yes), n (%) | 19 (43.2) | 24 (68.6) | 0.02<sup>¶</sup> |
| Antiseizure Medication Use (yes), n (%) | 28 (63.6) | — | — |
| *Oxcarbazepine* | 14 (31.8) |  |  |
| *Levetiracetam* | 13 (29.5) |  |  |
| *Lacosamide* | 1 (2.3) |  |  |
| Spike-wave laterality, n (%) |  |  |  |
| *Unilateral* | 24 (54.5) | — |  |
| *Bilateral* | 20 (45.5) | — | — |

§ Two tailed unpaired t-test
¶ Wald Chi-Square test

### 3.3 Thalamic CBF

#### 3.3.1 Group differences in thalamic CBF (Table 2, Figure 2)

Children with SeLECTS demonstrated elevated thalamic CBF (52.7 ± 1.5 mL blood / 100 g tissue min) compared to controls (46.9 ± 1.7 mL blood / 100 g tissue min; p = 0.014; Model 1). Thalamic CBF increased with age, and after adjusting for age, children with SeLECTS maintained higher thalamic CBF than controls (Model 2). The rate of age-related CBF increase did not differ between groups (Model 2a). Thalamic CBF did not significantly differ by sex, and after adjusting for sex, children with SeLECTS continued to show higher thalamic CBF than controls (Model 3). Neither age nor sex showed significant interactions with group, so these interaction terms were omitted from subsequent models. When adjusting for both age and sex, children with SeLECTS (51.9 ± 1.5 mL blood / 100 g tissue min) had higher thalamic CBF than controls (47.4 ± 1.7 mL blood / 100 g tissue min; p = 0.049; Model 4).

**Table 2.**
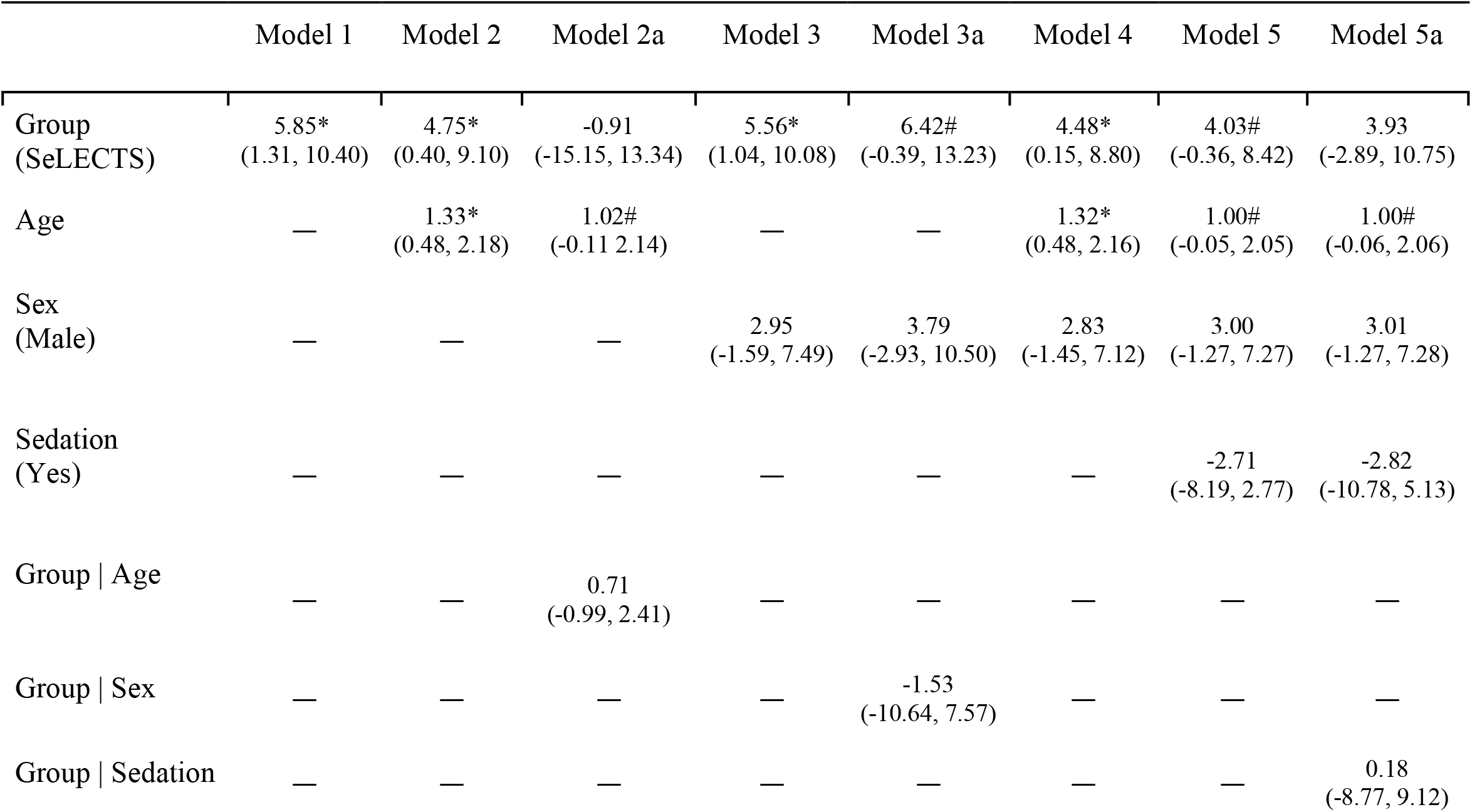
Linear mixed models of thalamic CBF in children with SeLECTS (n = 44) and controls (n = 35). Unstandardized coefficients (B) are reported with 95% confidence intervals in parentheses. * p<0.05; # p<0.10

**Figure 2.**
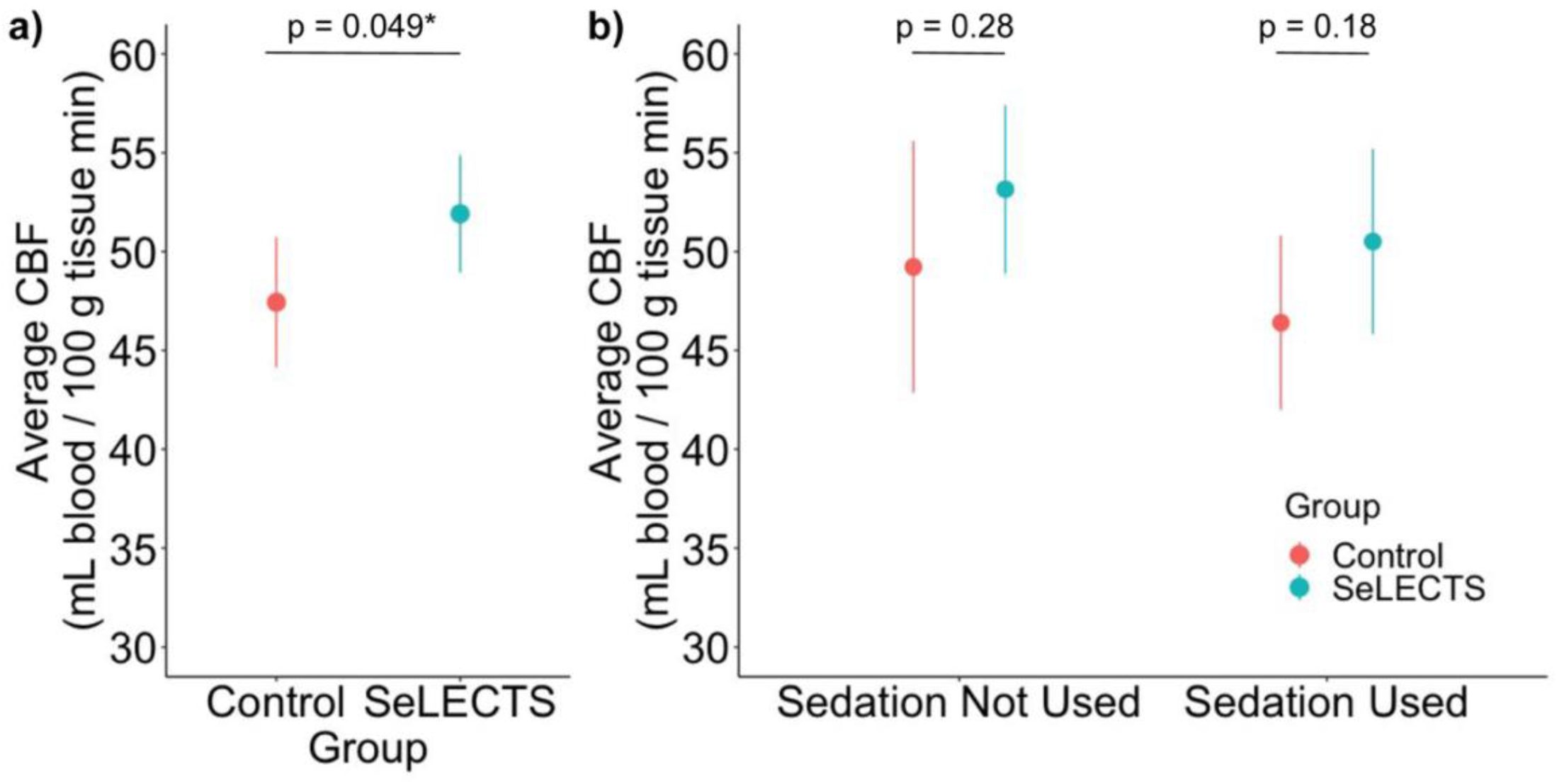
Group Differences in Thalamic Blood Flow. Plots show estimated marginal mean CBF values with 95% confidence intervals, adjusted for age and sex. **a)** Children with SeLECTS have higher thalamic blood flow than controls. **b)** Effect sizes remain consistent when stratified by sedation status, with similar magnitude differences observed in both sedated and unsedated children, supporting the robustness of the overall group difference despite reduced statistical power in these smaller subgroups.

In our sensitivity analysis including sedation status, the group difference became marginally significant (p = 0.08; Model 5). However, the magnitude of CBF differences between SeLECTS and controls was consistent in both sedated (4.1 ± 3.0 mL blood / 100 g tissue min) and unsedated (3.9 ± 3.6 mL blood / 100 g tissue min) children, with no significant group-by-sedation interaction (Model 5a; Figure 2b). This consistency in effect sizes across sedation status supports the interpretation that observed differences reflect underlying pathophysiology rather than acute state-dependent changes.

#### 3.3.2 Spike Laterality & Thalamic Hemisphere (Table 3, Figure 3)

**Figure 3.**
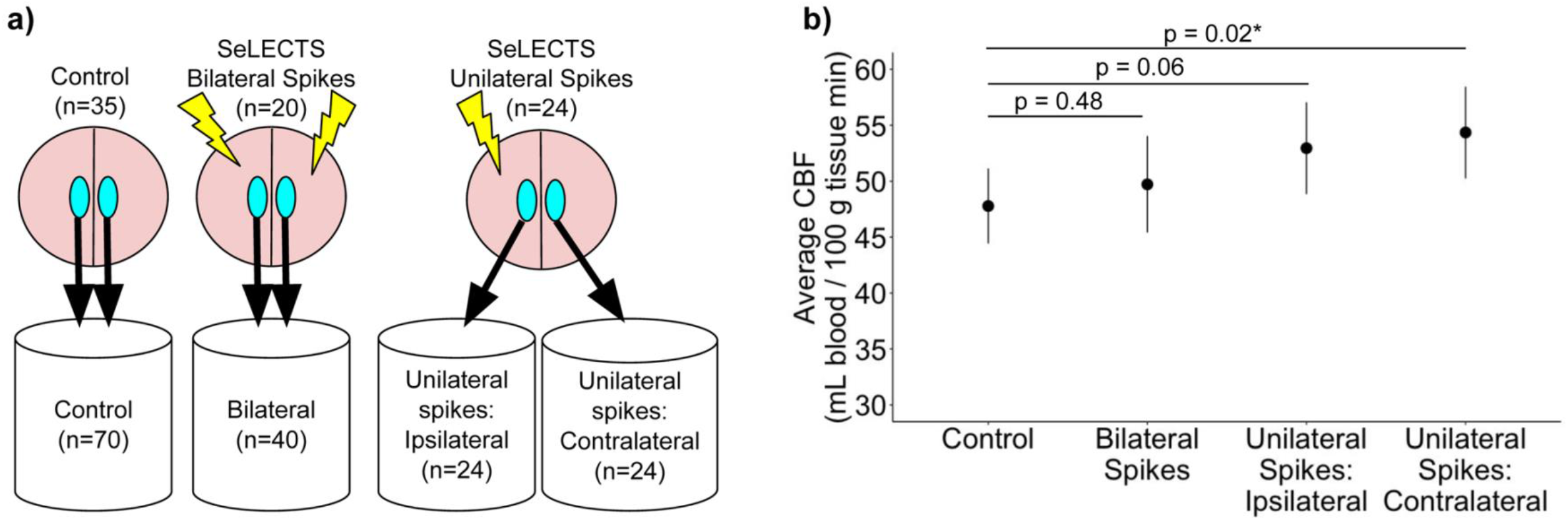
Thalamic blood flow relative to spikes. **a)** Method for classifying thalami by spike laterality. Blue ovals represent thalami and lightning bolts represent spikes. **b)** Plots show the estimated marginal mean CBF values with 95% confidence intervals for each of the different thalamic categories, adjusted for age, sex, and sedation during MRI. Children with bilateral spikes have a non-significant increase in CBF compared to controls, whereas children with unilateral spikes show a larger difference from controls.

Since left (46.7 ± 9.9 mL blood / 100 g tissue min) and right (47.0 ± 10.5 mL blood / 100 g tissue min) thalamic CBF did not differ in controls (p = 0.64), we grouped both hemispheres together, yielding 70 control thalamic measurements. Twenty children had bilateral spikes, yielding 40 bilateral CBF measurements. Twenty-four children had unilateral spikes, 12 with right hemisphere and 12 with left hemisphere, yielding 24 ipsilateral and 24 contralateral values.

**Table 3.**
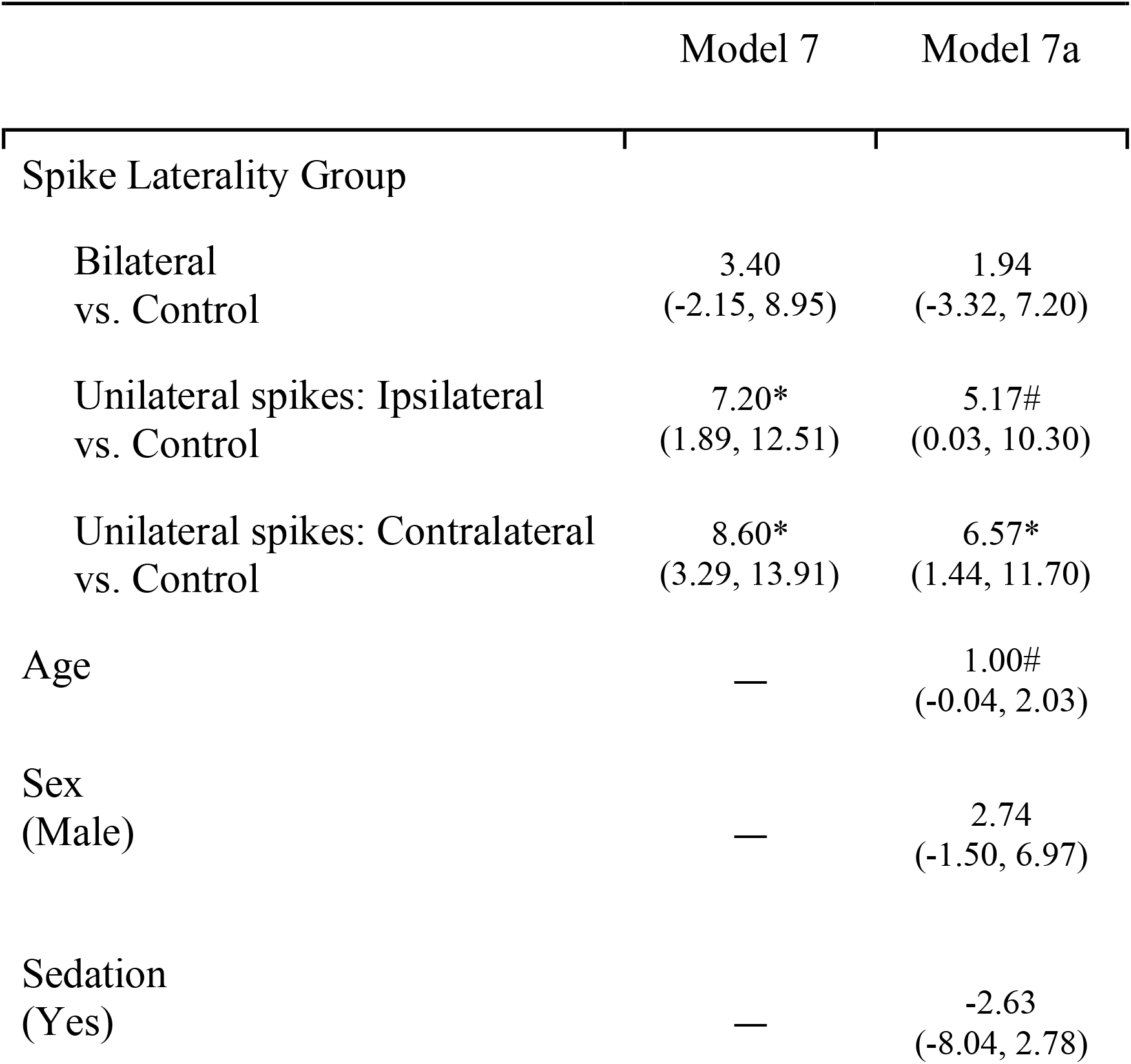
Linear mixed models of the effect of spike laterality on thalamic CBF. Unstandardized coefficients (B) are reported with 95% confidence intervals in parentheses. *p<0.05; # p<0.10

The most pronounced thalamic CBF elevations were observed in children with unilateral spikes. In unadjusted analyses, CBF in children with bilateral spikes was higher than that of controls, but the difference was not significant. In contrast, CBF in children with unilateral spikes was higher than controls in both ipsilateral and contralateral thalamic hemispheres, with the biggest difference surprisingly seen in the hemisphere contralateral to spikes (Model 7). After adjusting for age, sex and sedation (Model 7a), CBF in the thalamus contralateral to spikes (54.4 ± 2.1 mL blood / 100 g tissue min) was significantly higher than controls (47.8 ± 1.7 mL blood / 100 g tissue min, p = 0.017), and CBF in the thalamus ipsilateral (52.9 ± 2.0 mL blood / 100 g tissue min) was marginally elevated (p = 0.059) compared to controls; thalamic CBF in children with bilateral spikes (49.7 ± 2.2 mL blood / 100 g tissue min) was similar to controls (p = 0.48).

#### 3.3.3 ASMs (Table 4, Figure 4)

Of the 44 children with SeLECTS, 28 were on ASMs (oxcarbazepine n = 14, levetiracetam n = 13, lacosamide n = 1). Because only one child was on lacosamide, we excluded this child from analysis. In our unadjusted model (Model 8), elevated thalamic CBF relative to controls was seen in children with SeLECTS not on medication (p = 0.030) and children with SeLECTS on oxcarbazepine (p = 0.006). After adjusting for age, sex, and sedation (Model 8a), children taking oxcarbazepine demonstrated significantly elevated CBF (54.6 ± 2.6 mL blood / 100 g tissue min; p = 0.032) compared to controls (47.7 ± 1.7 mL blood / 100 g tissue min). Children taking levetiracetam had CBF values (48.3 ± 2.7 mL blood / 100 g tissue min; p = 0.85) indistinguishable from controls. Unmedicated children showed intermediate CBF values (52.7 ± 2.5 mL blood / 100 g tissue min; p = 0.11) that, while not statistically significant, were numerically similar to the oxcarbazepine group.

**Table 4.**
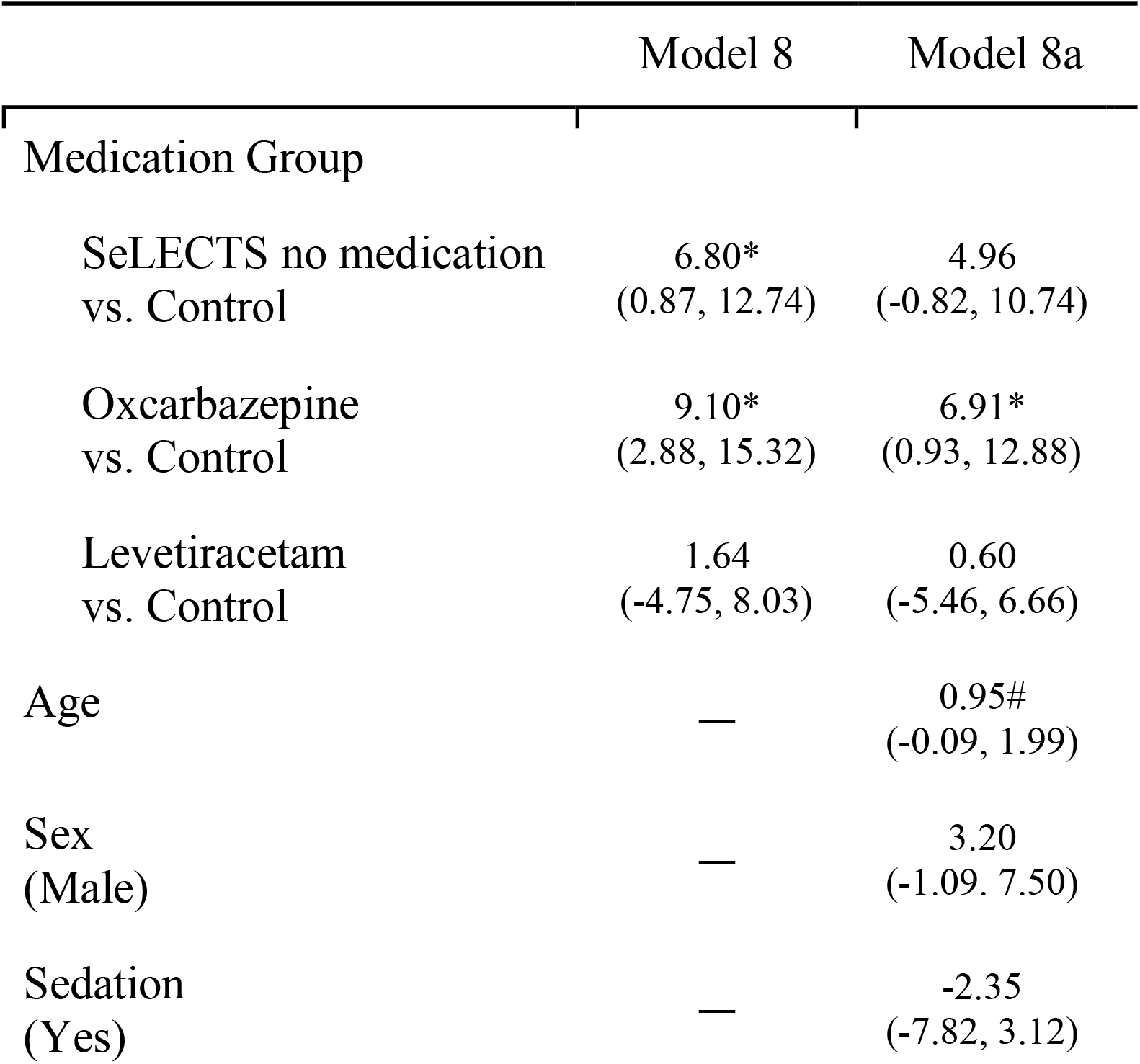
Linear mixed models of the effect of antiseizure medication on thalamic CBF. Unstandardized coefficients (B) are reported with 95% confidence intervals in parentheses. *p<0.05; # p<0.10

**Figure 4.**
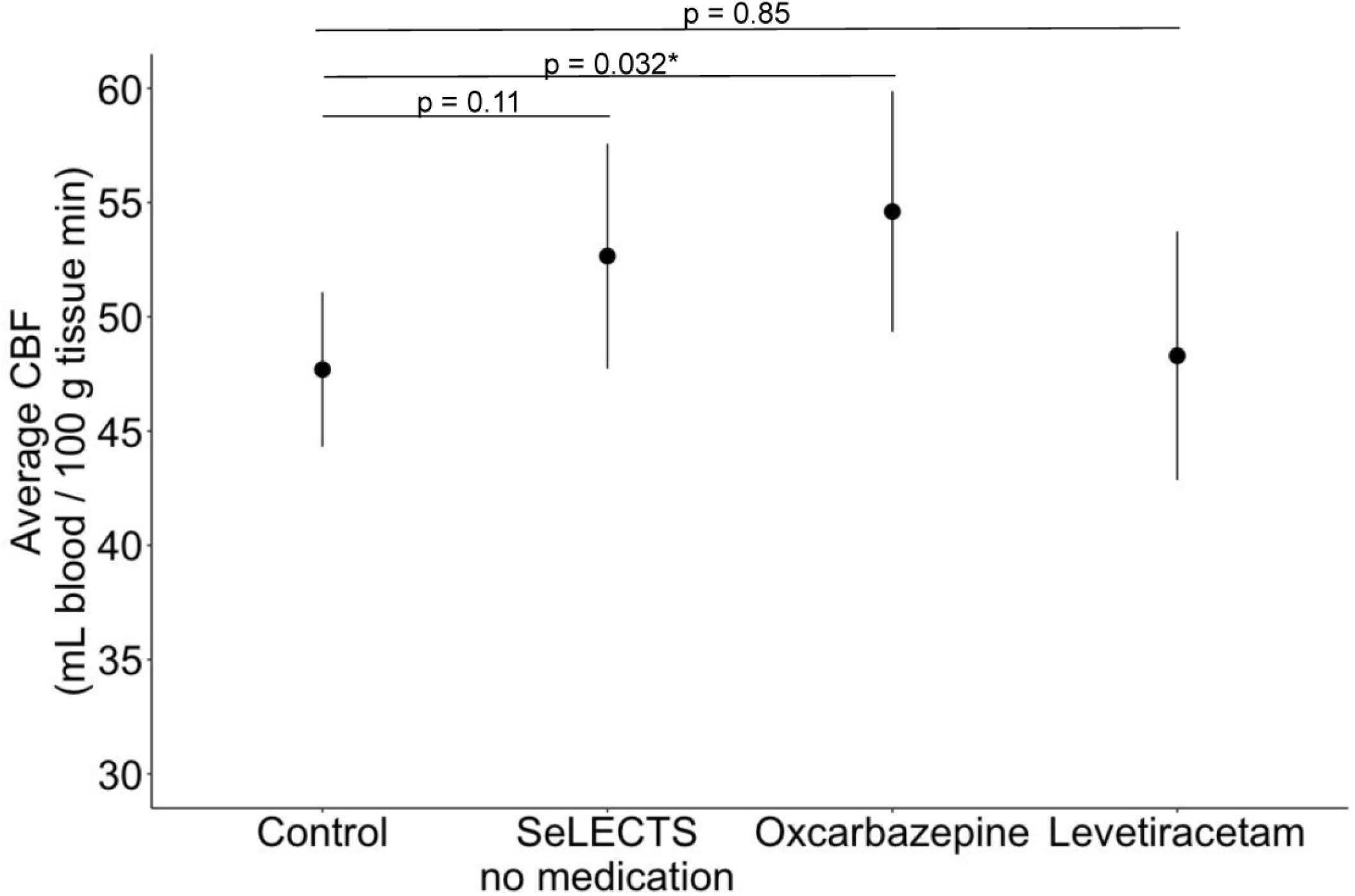
Thalamic blood flow by anti-seizure medication type. Plots show estimated marginal means with 95% confidence intervals, adjusted for age, sex, and sedation during MRI. Children with SeLECTS on oxcarbazepine have a significantly higher thalamic CBF than controls.

Our study was not powered to assess the effect of both spike laterality and medication use in the same model. Since oxcarbazepine tends to be prescribed for clearly focal epilepsies, whereas levetiracetam is used for both focal and generalized ones, we assessed whether those with unilateral spikes were more likely to be prescribed oxcarbazepine. Proportion of children with unilateral spikes was similar across medication groups (unmedicated: 8/16, 50%; oxcarbazepine: 8/14, 57%; levetiracetam: 8/13, 62%; Chi-square p = 0.82), suggesting independent effects of spike laterality and medication on thalamic CBF.

## 4.0 Discussion

The thalamus is increasingly recognized as being involved in SeLECTS pathophysiology. We compared thalamic blood flow, a marker of metabolism, in consecutively scanned children with SeLECTS to that in clinical controls. We find that children with SeLECTS have higher thalamic CBF than controls when compared as an overall group, though this difference became marginally significant after adjusting for sedation status at time of MRI. The most compelling insights emerged from subgroup analyses, which demonstrated that children with unilateral spikes and those receiving oxcarbazepine exhibit the most pronounced alterations in thalamic perfusion patterns.

The overall elevation in thalamic CBF observed in children with SeLECTS aligns with emerging evidence that the thalamus plays a central role in this epilepsy syndrome. Previous neuroimaging studies have documented altered structural and functional connectivity between the thalamus and sensorimotor cortices (Thorn et al., 2020; Kwon et al., 2022), and electrophysiological investigations have revealed that spike frequency inversely correlates with sleep spindle frequency, suggesting disruption of thalamocortical sleep circuits (Kramer et al., 2021). Our findings extend these observations by providing evidence that thalamic function, as measured by CBF, appears to be altered in SeLECTS, though the overall group comparison became marginally significant after controlling for potential confounders. The preservation of similar effect sizes in both sedated and unsedated children (4.1 vs 3.9 mL blood / 100 g tissue min) suggests that this marginal significance likely reflects our limited sample size rather than absence of a true pathophysiological difference, and also supports the interpretation that elevated thalamic blood flow is a consistent feature in SeLECTS, present regardless of behavioral state. Additionally, the fact that propofol, which suppresses epileptiform discharges (Meyer et al., 2006) and reduces thalamic CBF in adults (Saxena et al., 2019), did not eliminate group differences provides further evidence that thalamic hyperperfusion represents an intrinsic characteristic rather than an acute consequence of ongoing spike activity.

The differential effects of ASMs on thalamic CBF provide critical insights into SeLECTS pathophysiology and treatment mechanisms. Children taking oxcarbazepine demonstrated the highest thalamic CBF, unmedicated children showed intermediate elevations, and those taking levetiracetam had lower CBF which was similar to controls. This pattern may reflect differential effects of these medications on spike activity, as oxcarbazepine can paradoxically activate centrotemporal spikes in some children (Vendrame et al., 2007) while levetiracetam suppresses spikes (Kanemura et al., 2018; Zhang et al., 2018). The intermediate CBF observed in unmedicated children, while not achieving statistical significance (p=0.11), were numerically closer to the oxcarbazepine group than to controls, supporting the hypothesis that elevated thalamic blood flow is intrinsic to SeLECTS pathophysiology and that levetiracetam may normalize this underlying hyperperfusion whereas oxcarbazepine fails to suppress—and potentially exacerbates—this hyperperfusion.

The observation that children with unilateral spikes demonstrated the highest thalamic CBF values, particularly in the hemisphere contralateral to spike activity, represents a novel and unexpected finding that warrants careful consideration. This contralateral hyperperfusion pattern may reflect several potential mechanisms. The thalamus serves critical functions in state regulation and sensorimotor integration (Shine et al., 2023). In the context of SeLECTS, where normal thalamocortical oscillations are disrupted (Kramer et al., 2021), the contralateral thalamus may exhibit compensatory hyperactivity to maintain normal network function. This increased metabolic demand could manifest as elevated CBF in an attempt to preserve sleep architecture or maintain normal sensorimotor function. Alternatively, this pattern could represent an interictal phenomenon analogous to observations in other epilepsy syndromes, where epileptogenic regions typically show decreased metabolism during interictal periods (Chassoux et al., 2004). In temporal lobe epilepsy, the ipsilateral thalamus demonstrates reduced perfusion during interictal states as measured by single-photon emission computed tomography (Yune et al., 1998), suggesting that baseline thalamic hyperperfusion in SeLECTS may be asymmetrically suppressed in the hemisphere generating spikes. The intermediate CBF observed in children with bilateral spikes, falling between unilateral spike patients and controls, may indicate bilateral suppression of thalamic blood flow or represent a distinct pathophysiological subtype.

These medication-related findings contrast with established effects of these medications on brain metabolism documented in other epilepsy syndromes. Sodium channel blockers generally reduce cerebral metabolism, with FDG-PET studies demonstrating significant decreases in thalamic glucose metabolism following lamotrigine initiation in adults with idiopathic generalized epilepsy (Joo et al., 2006) and similar trends with carbamazepine in adults with focal epilepsy (Theodore et al., 1989). Levetiracetam has been less well studied, with one small FDG-PET study showing increased metabolism in certain brain regions only in patients who achieved seizure control (Kim et al., 2015). Given these established metabolic suppression effects of sodium channel blockers, the elevation of thalamic CBF observed with oxcarbazepine treatment in our SeLECTS cohort may suggest that thalamic hypermetabolism is a unique part of SeLECTS pathophysiology, differing from other epilepsy syndromes studied previously. Alternatively, and more likely, these discrepancies may reflect methodological differences between CBF measurements and FDG-PET, as studies have demonstrated variable concordance between these modalities in both healthy individuals and patients with epilepsy (Cha et al., 2013; Boscolo Galazzo et al., 2016; Gaillard et al., 1995). Supporting this second explanation, an FDG-PET study in D/EE-SWAS, a condition closely related to SeLECTS with frequent sleep-potentiated IEDs, found decreased thalamic metabolism (Agarwal et al., 2016). While the exact direction of metabolic changes remains unclear, our findings demonstrate that CBF provides a non-invasive, radiation-free approach to investigating thalamic function in pediatric epilepsy populations, offering opportunities for repeated measurements and longitudinal studies that could clarify these mechanistic questions.

### Limitations

Several limitations merit consideration. First, our clinical controls may not fully represent neurotypical children, despite systematic exclusion of children with disorders known to affect CBF. This limitation reflects the constraints of retrospective pediatric neuroimaging research and underscores the need for prospective studies with healthy controls. Second, selection bias may influence our findings, as neuroimaging is not routinely performed in all children with SeLECTS, and clinicians may preferentially image children with atypical presentations, frequent seizures, or unilateral spike patterns, potentially limiting the generalizability of our findings to the broader SeLECTS population. However, the proportion of unmedicated children in our cohort (36%) aligns with epidemiological data from the broader SeLECTS population (Lacey et al., 2024), suggesting reasonable generalizability. Third, the temporal disconnect between electroencephalographic recordings and magnetic resonance imaging acquisition prevents direct correlation of spike burden with CBF measurements. Multiple factors including behavioral state (Berroya et al., 2005), sedation (Meyer et al., 2006), and ASMs (Kanemura et al., 2018; Vendrame et al., 2007) can alter spike frequency, limiting our ability to establish causal relationships between epileptiform activity and thalamic perfusion. Additionally, spike laterality can evolve over time (Tenney et al., 2016), potentially introducing misclassification bias. Finally, our sample size constrained statistical power for complex multivariable analyses and prevented simultaneous examination of spike laterality and medication effects within the same model.

### Clinical Relevance & Future Direction

Our findings contribute to a growing understanding of how ASMs affect neurophysiology in SeLECTS, which may inform treatment strategies. The differential effects of levetiracetam and oxcarbazepine on thalamic blood flow align with emerging clinical evidence suggesting that levetiracetam may be superior for both seizure control and cognitive outcomes (Asadi-Pooya et al., 2019; Cheng et al., 2022). Our neurobiological findings provide a potential mechanistic basis for these clinical observations, suggesting that normalization of thalamic hyperperfusion may contribute to improved outcomes. More broadly, these findings highlight the potential of thalamic blood flow measurements to illuminate disease mechanisms that remain poorly understood in epilepsy. Future studies could leverage this approach to test whether thalamic CBF patterns correlate with key clinical outcomes such as seizure control, cognitive function, or degree of spike suppression. Such investigations could help determine whether thalamic dysfunction is a driver of clinical symptoms, and whether interventions targeting thalamic metabolism could improve patient outcomes. Concurrent EEG-MRI studies could establish direct relationships between spike activity and thalamic perfusion, while longitudinal designs could track whether CBF changes predict clinical trajectories. Advanced imaging techniques enabling assessment of individual thalamic nuclei (Takahashi et al., 2022) may provide more precise understanding of which specific thalamic circuits are involved in SeLECTS pathophysiology.

## Author contributions: CRediT

Niki Iasinovschi: Writing – original draft, Writing – review & editing, Visualization, Validation, Methodology, Formal analysis, Data curation, Conceptualization.

Elizabeth Tong: Writing – review & editing, Validation, Software, Methodology, Data curation, Conceptualization.

Lucia Bicknell: Writing – review & editing, Validation, Methodology, Data curation, Conceptualization.

Fiona M. Baumer: Writing – original draft, Writing – review & editing, Validation, Project administration, Methodology, Funding acquisition, Formal analysis, Data curation, Conceptualization.

## Acknowledgements & Funding Sources

This work was supported by National Institute of Neurological Disorders and Stroke (K23NS116110) (FMB), the Stanford University Department of Human Biology (NI, LB) and Office of the Vice Provost for Undergraduate Education (NI), and a gift from the Principe & O’Farrell Family (FMB). We would like to acknowledge Dr. Virginia Marchman for assistance with statistical analyses, Beattie Goad who curated the SeLECTS dataset, and Dr. Kristen Yeom and the Frankovich Laboratory who contributed to the control dataset.

## Data Statement

The data used for this study contains clinical identifiers and thus cannot be made publicly available.

## Declaration of generative AI in scientific writing

During the preparation of this work the author(s) used Claude in order to revise the completed manuscript for improved readability. After using this tool/service, the authors reviewed and edited the content completely and take full responsibility for the content of the published article.

## Notes

### Competing Interest Statement

The authors have declared no competing interest.

